# Capitalizing on paradoxical activation of the MAPK pathway for treatment of Imatinib-resistant mast cell leukemia

**DOI:** 10.1101/2020.08.31.266734

**Authors:** Thomas Wilhelm, Marcelo A. S. Toledo, Ilka Simons, Christian Stuth, Vrinda Mohta, Ronja Mülfarth, Marcus Nitsche, Karin Maschke-Neuß, Susanne Schmitz, Anne Kaiser, Michel Arock, Martin Zenke, Michael Huber

**Affiliations:** Institute of Biochemistry and Molecular Immunology, Medical Faculty, RWTH Aachen University, Pauwelsstraße 30, 52074 Aachen, Germany; Institute for Biomedical Engineering, Department of Cell Biology, RWTH Aachen University Medical School, Aachen, Germany.; Helmholtz-Institute for Biomedical Engineering, RWTH Aachen University, Aachen, Germany.; Department of Hematology, Oncology, Hemostaseology and Stem Cell Transplantation, Faculty of Medicine, RWTH Aachen University Medical School, Aachen, Germany.; Department of Hematological Biology, Pitié-Salpêtrière Hospital, Pierre et Marie Curie University (UPMC), 83 Boulevard de l’Hôpital, 75013 Paris, France.

**Author notes:** Corresponding author: Prof. Dr. Michael Huber Institute of Biochemistry and Molecular Immunology RWTH Aachen University Pauwelsstr. 30, 52074 Aachen, Germany.

## Abstract

Prevention of fatal side effects during cancer therapy of cancer patients with high-dosed pharmacological inhibitors is to date a major challenge. Moreover, the development of drug resistance poses severe problems for the treatment of patients with leukemia or solid tumors. Particularly drug-mediated dimerization of RAF kinases can be the cause of acquired resistance, also called “paradoxical activation”. Here we re-analyzing the effects of different tyrosine kinase inhibitors (TKIs) on the proliferation, metabolic activity, and survival of the Imatinib-resistant, KIT^V560G,D816V^-expressing human mast cell (MC) leukemia (MCL) cell line HMC-1.2. We observed that low concentrations of the TKIs Nilotinib and Ponatinib resulted in enhanced proliferation, suggesting paradoxical activation of the MAPK pathway. Indeed, these TKIs caused BRAF-CRAF dimerization, resulting in ERK1/2 activation. The combination of Ponatinib with the MEK inhibitor Trametinib, at nanomolar concentrations, effectively suppressed HMC-1.2 proliferation, metabolic activity, and induced apoptotic cell death. Effectiveness of this drug combination was recapitulated in the human *KIT* D816V MC line ROSA KIT^D816V^ and in *KIT* D816V hematopoietic progenitors obtained from in patient-derived induced pluripotent stem cells (iPS cells). In conclusion, mutated KIT-driven Imatinib resistance can be efficiently bypassed by a low concentration combination of the TKI Ponatinib and the MEK inhibitor Trametinib, potentially reducing the negative side effects associated to MCL therapy.

## Introduction

The successful treatment of human malignancies caused by constitutively active tyrosine kinases using tyrosine kinase inhibitors (TKIs) is one of the major breakthroughs in cancer therapy. The first and still best example for a successful TKI is Imatinib (initially named CGP 57148; a.k.a GLEEVEC^→^ or STI 571), which was described first in 1998 (Jonuleit et al. 1998) and over time has replaced allogeneic stem cell transplantation in the therapy of *BCR-ABL1*-positive chronic myelogenous leukemia (CML) (Druker 2008). Imatinib also effectively inhibits the wild-type receptor tyrosine kinase KIT (CD117) and some of its mutants and hence is useful, for instance, in therapies of gastrointestinal stromal tumors (Roskoski 2018). Although Imatinib is able to inhibit most activating point mutants of BCR-ABL1 and KIT, mutants completely resistant to Imatinib exist (e.g. KIT D816V and BCR-ABL1T315I) and aggravate therapy of respective patients. The situation is particularly detrimental in patients with systemic mastocytosis (SM), who express *KIT* D816V in more than 80% of all cases (Valent et al. 2017).

SM is a heterogeneous mast cell (MC) disorder characterized by abnormal MC infiltration into different organs and tissues, and increased release of MC mediators. SM can span from an indolent form to forms with poor prognosis, namely aggressive SM and MC leukemia (MCL) (Valent et al. 2017). As mentioned, KIT D816V, which is expressed in the majority of all SM cases, is resistant to the first generation TKI Imatinib and only weakly responsive to the second generation of TKIs e.g. Nilotinib (AMN107) (Gleixner 2006). Nevertheless, some of the third generation TKIs like Ponatinib and Midostaurin are able to inhibit KIT D816V (Gleixner et al. 2013). Unfortunately, due to the low selectivity of these and other TKIs, harmful side effects can occur. For instance, it has been shown that both Nilotinib and Ponatinib can cause cardiac and vascular pathologies (Isfort & Brümmendorf 2018; Gambacorti-Passerini et al. 2016), whereas treatment with Midostaurin induced serious GI-tract side effects and frequent hematologic adverse events (DeAngelo et al. 2018).

Activation of KIT by binding of its natural ligand, stem cell factor (SCF), or by activating mutations results in the induction of several signaling pathways in control of cell proliferation, survival, and metabolism, such as the mitogen-activated protein kinase (MAPK) pathway and the phosphatidylinositol-3-kinase (PI3K) pathway, as well as activation of the transcription factor STAT5 (Buet et al. 2012). KIT-mediated activation of the MAPK pathway is initiated by autophosphorylation of several tyrosine residues in KIT enabling the recruitment of the adaptor protein SHC, followed by binding of the GRB2/SOS complex. The guanine nucleotide exchange factor SOS is then responsible for the GDP-to-GTP exchange in RAS, with RAS-GTP mediating the dimerization and activation of kinases of the RAF family (e.g. BRAF and CRAF), resulting in the activation of the dual-specificity kinases MEK1/2. Finally, MEK1/2 phosphorylate and activate ERK1/2, which have cytosolic as well as nuclear targets and, in addition to activating functions, are involved in negative feedback regulation of the MAPK pathway. MAPK pathway activation has been identified in a wide range of malignancies, promoting proliferation and survival (Burotto et al. 2014).

In a high percentage of melanoma patients, the BRAF V600E mutant is expressed, which is active as a monomer in a RAS-GTP-independent manner (Davies et al. 2002). The BRAF V600E-selective inhibitor Vemurafenib was shown to trigger the MEK/ERK pathway in BRAF V600E-positive melanoma cells that had acquired an additional *RAS*-activating mutation (Poulikakos et al. 2010; Callahan et al. 2012). In such a situation, Vemurafenib mediated heterodimerization of BRAF V600E with non-mutated CRAF in an active RAS-dependent manner. This then allowed for MEK activation *via* the still activatable CRAF molecule in the BRAF V600E-CRAF heterodimer. This phenomenon was called “paradoxical activation”. Meanwhile, different TKIs (e.g. Imatinib, Nilotinib, and Dasatinib) have been demonstrated to unexpectedly interact with RAF proteins thereby causing their dimerization and unconventional activation by active RAS, resulting in the increased stimulation of the MEK/ERK pathway (Packer et al. 2011; Poulikakos et al. 2010; Hatzivassiliou et al. 2010). Hence, an alternative to overcome unwanted TKI-mediated MAPK pathway activation is the therapeutic intervention with MEK inhibitors, which are widely used for cancer treatment in *in vitro* studies as well as in clinical trials (reviewed by Caunt (Caunt et al. 2015)).

Allosteric MEK inhibitors, such as PD0325901 or PD184352 (CI-1040) target a unique inhibitor-binding pocket that is separated from, but adjacent to, the Mg^2+^-ATP-binding site in MEK1/2 (Ohren et al. 2004), stabilizing their inactive conformation. However, such allosteric MEK inhibitors prevent negative feedback phosphorylation of BRAF and CRAF by ERK1/2, resulting in accumulation of inhibited, but still phosphorylated MEK1/2. A reduction of the cellular MEK inhibitor/MEK molecule ratio, provoked by cellular resistance mechanisms or missing patient compliance, can then cause vehement re-activation of the MEK/ERK pathway (Caunt et al. 2015). MEK inhibitors of a newer generation, like Trametinib (GSK 1120212; JTP-74057), alter the conformation of the activation loop of MEK, thereby preventing both MEK phosphorylation by RAF kinases and catalytic activity of MEK (Lito et al. 2014).

In the present study, we have re-analyzed the consequences of TKI treatment on human *KIT* D816V-positive MCs and *KIT* V560G,D816V-positive MCL cells. We were able to demonstrate that TKI concentration insufficient to inhibit cell proliferation are able to induce paradoxical RAF activation characterized by enhanced MEK activation and cell proliferation. The combination of low-dosed TKIs with MEK inhibitors was able to synergistically induce cell death. A comparable mechanism was demonstrated in patient-specific iPS cell-derived hematopoietic cells, despite heterogeneous expression of additional mutations. In conclusion, we present a way to enhance the efficacy of TKI treatment in MCL to potentially reduce negative side-effects by allowing use of lower TKI concentrations resulting in improved patients outcome.

## Material and Methods

### Cell culture

HMC-1.2 (*KIT* V560G,D816V) MCL cells were kindly provided by Dr. J. Butterfield (Mayo Clinic, Rochester, MN) (Butterfield et al. 1988). They were maintained in RPMI-1640 medium (Gibco, Thermo Fisher Scientific) supplemented with 10% fetal bovine serum (FBS) and 10.000 units penicillin + 10 mg/ml streptomycin (all from Sigma-Aldrich) in an atmosphere containing 5% CO_2_. The medium was renewed twice a week.

ROSA KIT^D816V^ cells were cultured as previously described (Saleh et al. 2014) in IMDM medium supplemented with 10.000 units penicillin + 10 mg/ml streptomycin, 100 mM sodium pyruvate, MEM vitamin solution, MEM amino acids, 200 mM L-glutamine, insulin-transferrin-selenium (all from Gibco, Thermo Fisher Scientific), and 10 % FBS.

### iPS cell lines

The generation of the iPS cell lines used in the present work was described previously (Toledo et al. 2020). Further information on each iPS cell line can be found at the Human Pluripotent Stem Cell Registry (www.hpscreg.eu) where patient 1 control_1 and *KIT* D816V_1 iPS cell lines are referred to as UKAi004-A and UKAi004-C, respectively, and patient 2 control_1 and *KIT* D816V_1 iPS cell lines are referred to as UKAi008-B and UKAi008-C, respectively.

### Hematopoietic differentiation of iPS cells

The differentiation of iPS cells towards the hematopoietic lineage was performed through the formation of embryoid bodies (EB) by the spin-EB method modified from (Liu et al. 2015).

Of note, fromday 7 to 14 after EB formation, no BMP4 or VEGF was added to the medium and, from day 10 to 14, emerging hematopoietic cells and EBs were cultivated in serum free medium (SFM) supplemented only with 100 ng/ml recombinant human stem cell factor (rhSCF, Milteny Biotech) and 30 ng/ml recombinant human interleukin 3 (rh IL-3, both Peprotech).

From days 10 to 14 after EB formation, emerging hematopoietic cells and EBs were cultivated in SFM ((Liu et al. 2015) supplemented with recombinant human 10 ng fiblroblast growth factor type 2 (rhFGF-2, Peprotech), recombinant human 50 ng/mL rhSCF and 30 ng/mL rhIL-3. On day 14, hematopoietic cells were transferred to a 10 cm-dish and further cultivated for 14 days on SFM medium supplemented with 100 U/ml penicillin, 100 µg/ml streptomycin (Thermo Fisher Scientific),100 ng/ml rhSCF, 50 ng/ml recombinant human fms-related tyrosine kinase 3 ligand (rhFLT3L, Peprotech), 30 ng/ml rhIL-3 (all from Peprotech), and 10 ng/ml interleukin 6/soluble interleukin 6 receptor fusion protein (hyper-IL-6), (Fischer et al. 1997) at a cell density of 0.5 - 1.0 x 10^6^ cells/ml. Partial medium change was performed every 4 days. After 14 days of expansion, cells were harvested and starved for at least 1 hour in RPMI-1640 medium supplemented with 100 U/ml penicillin, 100 µg/ml streptomycin, 2 mM L-glutamine (all from Thermo Fisher Scientific). Cells were then enriched for KIT expressing cells by magnetic activation cell sorting (MACS) using CD117 MicroBead Kit (Miltenyi Biotec), following manufactureŕs instructions.

### Apoptosis assay

HMC-1.2 and ROSA KIT^D816V^ cells were seeded at a density of 3.5 x 10^5^ cells/ml and treated with the indicated substances for 72h. After treatment HMC-1.2 cells were incubated with Annexin V-Alexa Fluor 647 (Alexis Biochemicals) in culture medium for 20 min at 4°C in the dark. Immediately before analysis by flow cytometry, propidium iodide (1 μg/ml) was added and the cells were analyzed on a flow cytometer (Canto II; BD Biosciences). ROSA KIT^D816V^ cells were stained with Annexin V-Alexa Fluor 647 (Alexis Biochemicals) and 7AAD in culture medium for 20 min at 4°C in the dark.

### Reagents

PD0325901 was purchased from Axon Medchem and Imatinib, Nilotinib, Ponatinib and Trametinib were purchased from Selleckchem. DMSO was obtained from Carl Roth GmbH & Co.

### Western blotting, Immunoprecipitation, and antibodies

Pelleted cells were solubilized with 0.5% NP-40 and 0.5% sodium deoxycholate in 4°C phosphorylation solubilization buffer (Wilhelm et al. 2018). After normalizing for protein content, lysates were supplemented with Lämmli buffer, boiled for 5 minutes at 95°C and subjected to SDS-PAGE and subsequent Western blot analysis as previously described (Wilhelm et al. 2018). The following antibodies were used for detection of BRAF (F-7, sc-5284), CRAF (C-12, sc-133), KIT (M-14, sc-1494), STAT5a (A-7, sc-166479), PARP-1 (H-250, sc-7150), GAPDH (6C5, sc-32233) and were purchased from Santa Cruz, whereas antibodies used for detection of pKIT (Y719, #3391), ERK1/2 (L34F12, # 4696), pERK1/2 (Thr202/Tyr204, #4370), α-pSTAT5 (Tyr694, #9351), α-Caspase3 (#9662), cleaved Caspase3 (Asp175, #9664), α-pMEK (Ser221, #2338) were purchased from Cell Signaling Technology. For RAF immunoprecipitation, 10^6^ cells were treated with TKIs for 3 h in 10 ml medium. At the end of this treatment, cells were washed and lysed after centrifugation in 500 µl PSB buffer. The lysate was incubated on a rotator for 1 h at 4°C followed by centrifugation. The supernatant was collected and 15 µl of anti-CRAF antibody was added and incubated O/N on a rotator at 4°C. Then 50 µl of Protein-G-Sepharose-Bead-Suspension were added and the resulting mixture was incubated at a rotator for 2 h at 4°C. The beads were then collected by centrifugation, washed three times, and boiled at 95°C with 10 µl 2X Lämmli buffer for 5min.

### RNA preparation and quantitative RT-PCR

RNA preparation and qRT-PCR has been described previously (Wilhelm et al. 2018). Briefly, RNA from 3 x 10^6^ HMC-1.2 cells was extracted using RNeasy Plus Mini Kit (Qiagen) according to the manufacturer’s instructions. Total RNA (1 μg) was reverse transcribed using Random hexamers (Roche) and Omniscript Kit (Qiagen) according to the manufacturer’s instructions. qRT-PCR was performed on a Rotorgene (Qiagen) by using SYBR green reaction mix (Bioline #QT650-02). Expression was normalized to the housekeeper HPRT. The relative expression ratio including primer efficiencies was calculated by the Pfaffl method (Pfaffl 2001). Primer sequences and efficiency data were as follows: BCL2L1 fwd ACT CTT CCG GGA TGG GGT AA, rev AGG TAA GTG GCC ATC CAA GC, 1.98; CCND1 fwd GCC CTC GGT GTC CTA CTT C, rev AGG AAG CGG TCC AGG TAG TT, 2.08; HPRT fwd TGA CAC TGG CAA AAC AAT GCA, rev GGT CCT TTT CAC CAG CAA GCT, 2.03

### Proliferation assays

Cells were seeded at a density of 3.5 x 10^5^ cells/ml and treated with the indicated concentrations of the test substances; solvent (DMSO)-treated cells served as controls. Cells were resuspended completely every 24 h and 50 μl from each well was diluted in 10 ml PBS for automated multi-parameter cell counting using a Casy cell counter (Innovatis). Metabolic activity was measured using the XTT Cell Proliferation Kit II (XTT) (Roche). Cells were seeded in microplates at a density of 3.5 x 10^5^ cells/ml (suspension culture grade, 96 wells, flat bottom) in a final volume of 100 µl culture medium per well in a humidified atmosphere (37°C, 5% CO_2_) for 72h. After the incubation period, 50 µl of the XTT labeling mixture was added to each well (final XTT concentration 0.3 mg/ml). Incubation of the microplate was for 3 - 4 h in a humidified atmosphere (e.g., 37°C, 5% CO_2_).

Spectrophotometric absorbance of the samples was measured using a microplate reader. The wavelength used to measure absorbance of the formazan product of the XTT assay was 475nm and the reference wavelength was 650nm. Sample values at 475nm were subtracted with medium controls (blanked) resulting in delta blanked values. Total absorbance was calculated by subtraction of delta blanked values (475nm) with their reference values at 650nm. These absorbance values (A_475nm_-A_650nm_) are shown in the respective figures.

### Statistical analysis

Data were generated from independent experiments. The statistical analysis and graphing of the data were performed using GraphPad Prism 8.30 (GraphPad Software, San Diego, CA 92108). ANOVA tests and one sample t-test were performed as noted in the respective figure legends. P values were considered statistically significant according to the following: * < 0.05, ** < 0.01, and *** < 0.001; ns indicates no significance. The individual number of independent biological replicates per experiments is shown in the legends.

## Results

### 1. Suboptimal concentrations of tyrosine kinase inhibitors enhance proliferation of HMC-1.2 cells

In contrast to KIT^WT^ and some activation mutations of KIT e.g. KIT V560G, KIT D816V is resistant to Imatinib and has limited response to second generation TKIs e.g. Nilotinib (Gleixner et al. 2006). Although third generation TKIs e.g. Ponatinib (Lierman et al. 2012) are able to inhibit KIT D816V, major challenges remain such as acquired resistance and severe side-effects (Isfort & Brümmendorf 2018; Gambacorti-Passerini et al. 2016). In our present study, we first re-evaluated in HMC-1.2 cells, expressing KIT V560G and KIT D816V, the anti-leukemic efficiencies of the TKIs Imatinib, Nilotinib, and Ponatinib. Whereas Imatinib, even at 10 µM, did not significantly impact on proliferation (determined by cell counting), metabolic activity (measured by XTT assays), and survival (analyzed by annexin V/propidium iodide (AV/PI) staining) of HMC-1.2 cells (Fig. 1A, D, G), Nilotinib and Ponatinib clearly suppressed proliferation as well as metabolic activity (Fig. 1B, C, E, F), and promoted cell death (Fig. 1H, I). From the dose-response analysis performed, it was obvious that Ponatinib was more effective than Nilotinib by a factor of approximately 10. Notably, less effective concentrations of these TKIs (Nilotinib, 1 µM; Ponatinib, 0.1 µM) significantly increased proliferation of HMC-1.2 cells (Fig. 1B, C). It has been shown for other leukemic cells (e.g. CML and ALL cells) that certain TKIs are able to bind to RAF kinases, induce their hetero-dimerization, and enable paradoxical activation of the MAPK pathway in the presence of active RAS (Packer et al. 2011). To determine if such mechanism is also functional in HMC-1.2 cells, these cells were treated with the solvent DMSO or increasing concentrations of the TKIs Imatinib, Nilotinib, and Ponatinib, and TKI-induced dimerization of CRAF with BRAF was analyzed by anti-CRAF immunoprecipitation followed by BRAF-specific immunoblotting. Whereas no dimerization was observed in DMSO-treated cells, Nilotinib and Ponatinib induced strong coprecipitation of BRAF with anti-CRAF antibodies (Fig. 1J). Particularly at low concentrations of Nilotinib and Ponatinib, which did not diminish phosphorylation of KIT, phosphorylation of ERK1/2 was enhanced, correlating with increased proliferation under these conditions (Fig. 1B, 1C). Imatinib treatment, on the other hand, resulted in modest CRAF/BRAF dimerization and increased ERK1/2 phosphorylation (Fig. 1A, J). In conclusion, treatment of HMC-1.2 cells with suboptimal concentrations of Nilotinib or Ponatinib resulted in solid paradoxical activation of RAF kinases correlating with enhanced proliferation.

**Figure 1:**
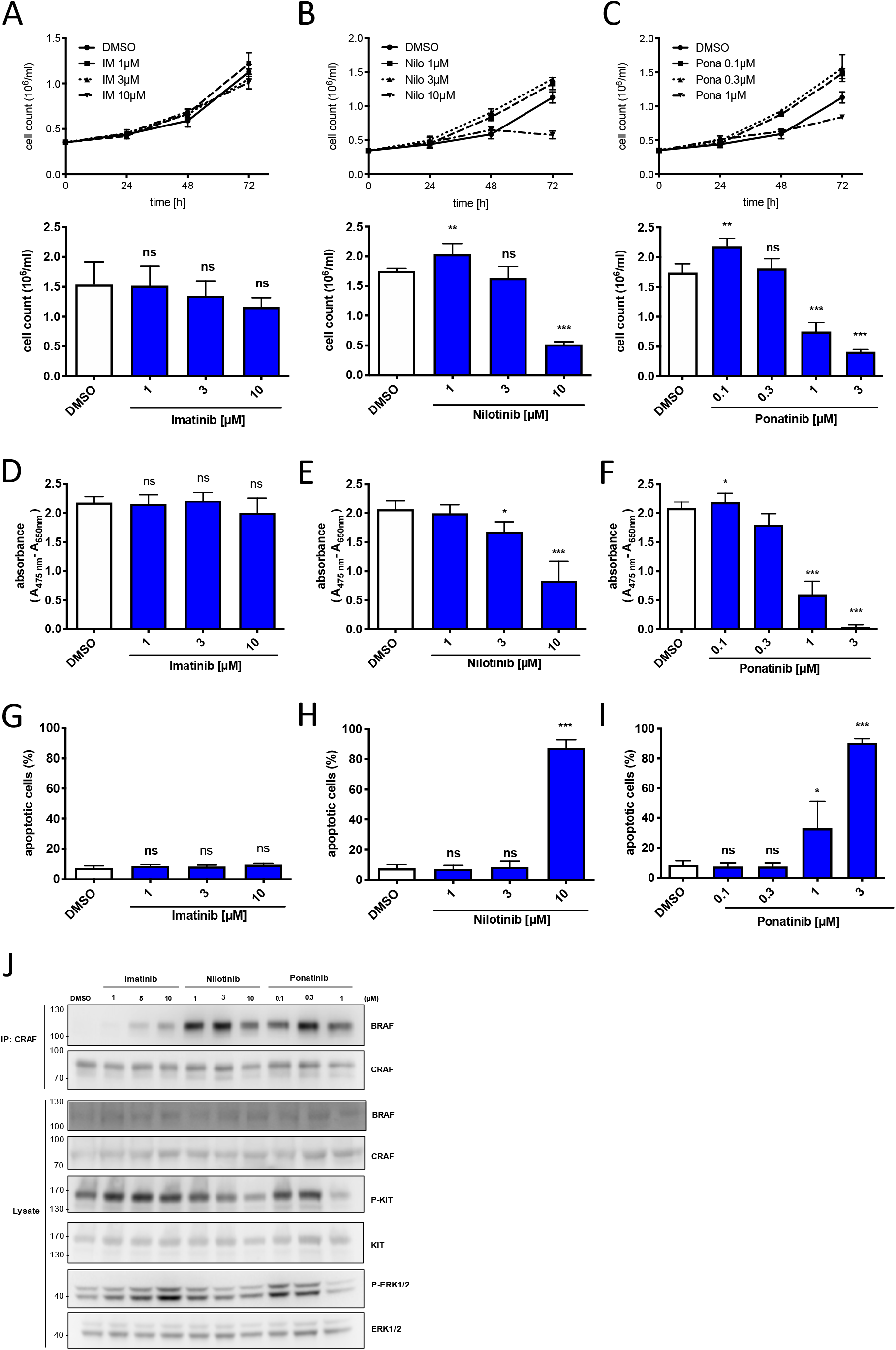
Tyrosine kinase inhibitors induce paradoxical RAF activation in HMC-1.2 cells. HMC-1.2 cells were seeded at a density of 3.5 x 10^5^ cells/ml and treated with the TKIs Imatinib, Nilotinib and Ponatinib at the indicated concentrations. The cell number of Imatinib (n=4) (A), Nilotinib (B) (n=4) or Ponatinib (C) treated cells was measured every 24 h for up to 72 h using a CASY Cell Counter (n=4). The upper panel shows a time course of an individual experiment. The lower panel shows the average of repetitive experiments (n=4) after 72 h. The metabolic activity of Imatinib (D) (n=3), Nilotinib (E) (n=4) and Ponatinib (F) (n=5) treated cells was analyzed after 72 h using XTT assays. Cell viability was measured after 72 h by FACS analysis using Annexin V and Propidium Iodide staining of Imatinib (G) (n=3), Nilotinib (H) (n=4) or Ponatinib (I) (n=4) treated cells. CRAF immunoprecipitations and total cell lysates from TKI treated cells treated for 3 h were analyzed by western blot for CRAF, BRAF and phospho-KIT, KIT (loading control), phosphor-ERK1/2, ERK1/2 (loading control) (J), representative experiment (n=4). Mean + SD, one-way ANOVA followed by Tukey (multiple comparison) post-test. * P < 0.05, ** P < 0.01, and *** P < 0.001; ns indicates no significance.

### 2. MEK inhibition exerts an anti-proliferative and pro-apoptotic effect in HMC-1.2 cells

Inhibition of the MEK-ERK pathway attenuates proliferation and survival of various cancer cells, in particular in several types of leukemias (reviewed by Steelman et al. 2011). Thus, we hypothesized that TKI-induced paradoxical activation of the MEK-ERK pathway could be counteracted by pharmacological inhibition of MEK leading to proliferation suppression and reduced survival of neoplastic cells. Therefore, we subjected HMC-1.2 cells to treatment with different concentrations of the selective MEK inhibitor PD0325901 to a) test if their proliferation and survival are dependent on the MEK-ERK pathway, and b) find out the lowest meaningful concentrations of PD0325901 to reduce potential side effects. The analysis of proliferation (Fig. 2A), metabolic activity (Fig. 2B), and survival (Fig. 2C) revealed a concentration-dependent suppression of these cellular traits by PD0325901. First apparent effects, though not yet significant, were observed upon treatment with 50-100 nM. As markers for proliferation and survival, we measured expression of the mRNAs of *CCND1* (coding for cyclin D1) and *BCL2L1* (coding for BCL_XL_), respectively. Indeed, 50 nM PD0325901 caused significant and strong reduction in expression of *CCND1* (Fig. 2D) and *BCL2L1* (Fig. 2E), substantiating the efficacy of PD0325901 in HMC-1.2 cells at this low concentration.

**Figure 2:**
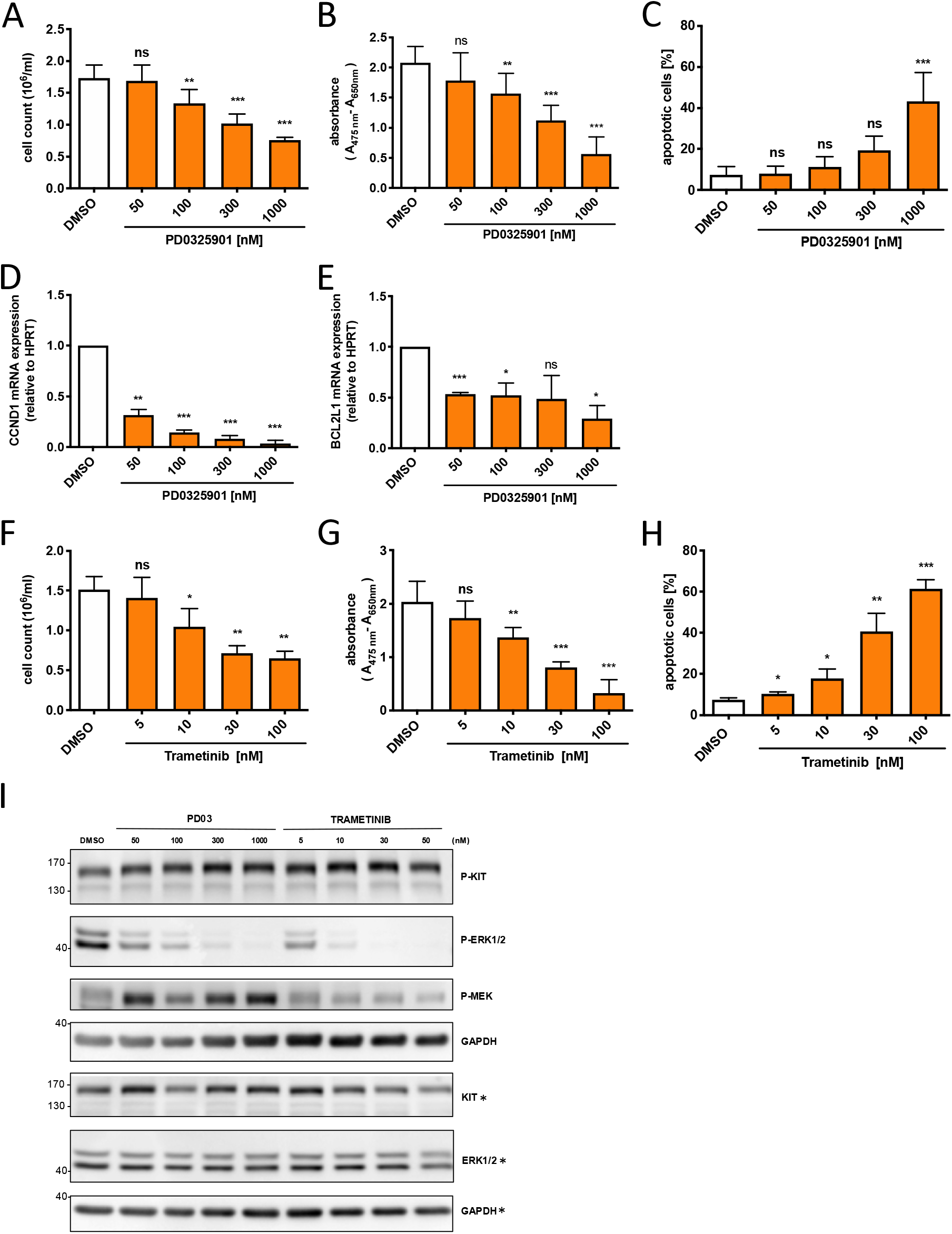
MEK inhibition by PD0325902 or Trametinib reduces proliferation and survival of HMC1.2 cells. HMC-1.2 cells were seeded at a density of 3.5 x 10^5^ cells/ml and treated with MEK inhibitors at the indicated concentrations for 72 h. After incubation of HMC-1.2 cells with PD0325901 for 72 h, Proliferation (A) (n=3) was measured using a Casy cell counter, metabolic activity was determined by XTT assay (B) (n=4) and survival was analyzed by Annexin V and Propidium Iodide staining (C) (n=3) using FACS. Expression of *CCND1* (n=3) (D) and *BCL2-L1* (n=3) (E) of PD0325901 treated HMC-1.2 cells for 3 h was measured by RT-qPCR. Proliferation (F) (n=4), metabolic activity (G) (n=3) or survival (H) (n=3) of Trametinib treated cells was analyzed after 72 h. Lysates of PD03 or Trametinib treated cells for 3 h were analyzed by western blot for phosphorylation of KIT, ERK1/2 and MEK. KIT, ERK1/2, and GAPDH served as loading controls (I), representative experiment (n=3). Mean + SD, one-way ANOVA followed by Tukey (multiple comparison) post-test. Figure D and E one sample t-test. * P < 0.05, ** P < 0.01, and *** P < 0.001; ns indicates no significance.

ERK as the final kinase in the MAPK cascade is able to negatively feedback onto upstream components of this pathway, such as MEK, RAF, and SOS (Shin et al. 2009). Hence, MEK inhibition by classical MEK inhibitors, such as PD0325901, causes upregulation of MEK phosphorylation in HMC-1.2 cells, whereas ERK1/2 is attenuated (Fig. 2I). A new class of MEK inhibitors, so-called “feedback busters”, is able to prevent accumulation of phosphorylated MEK after its inhibition. Trametinib, a “feedback buster” already used in the clinic, did suppress ERK phosphorylation in HMC-1.2 cells without increasing MEK phosphorylation (Fig. 2I). Moreover and importantly, Trametinib suppressed proliferation and metabolic activity as well as induced apoptosis of HMC-1.2 cells more effectively than PD0325901 (by a factor of 10) (Fig. 2 F-H).

### 3. Synergistic inhibition by PD0325901 and Nilotinib of growth and survival of HMC-1.2 cells

Paradoxical MAPK activation by TKI-mediated BRAF-CRAF complexes might sensitize HMC-1.2 cells to MEK inhibitors. Thus, we next addressed if the combination of low concentrations of the TKI Nilotinib and the MEK inhibitor PD0325901 would result in significant, synergistic suppression of proliferation and survival of HMC-1.2 cells. For this purpose, we decided to use 50 nM PD0325901 and 3 µM Nilotinib, since both inhibitors at the chosen concentrations did neither repress proliferation (Figs. 1B & 2A) nor reduce survival (Figs. 1H & 2C) of HMC-1.2 cells. Indeed, this combination of inhibitors resulted in a strong, synergistic suppression of proliferation (Fig. 3A, B) as well as metabolic activity (Fig. 3C), and was able to significantly reduce survival (Fig. 3D) of HMC-1.2 cells. To characterize the observed cell death more closely, we treated HMC-1.2 cells for 48 h with 50 nM PD0325901, two concentrations of Nilotinib (1 µM and 3 µM) as well as the respective combinations. Combined treatments caused cleavage of Caspase-3 and its target PARP1 (Fig. 3E), indicating Caspase-3 activation and induction of apoptosis. Of note, also single treatments resulted in marginal cleavage of Caspase-3 and PARP1, which, however, did not have a significant impact on survival of HMC-1.2 cells (Fig. 3D). Moreover, constitutive phosphorylation of the pro-survival transcription factor STAT5 was very sensitive to Nilotinib treatment, although the effect on KIT phosphorylation was marginal (Fig. 3E). Analysis of ERK1/2 phosphorylation proves the efficiency of PD0325901 (Fig. 3E). In conclusion, we demonstrated synthetic lethality by low concentrations of Nilotinib and PD0325901 in HMC-1.2 cells.

**Figure 3:**
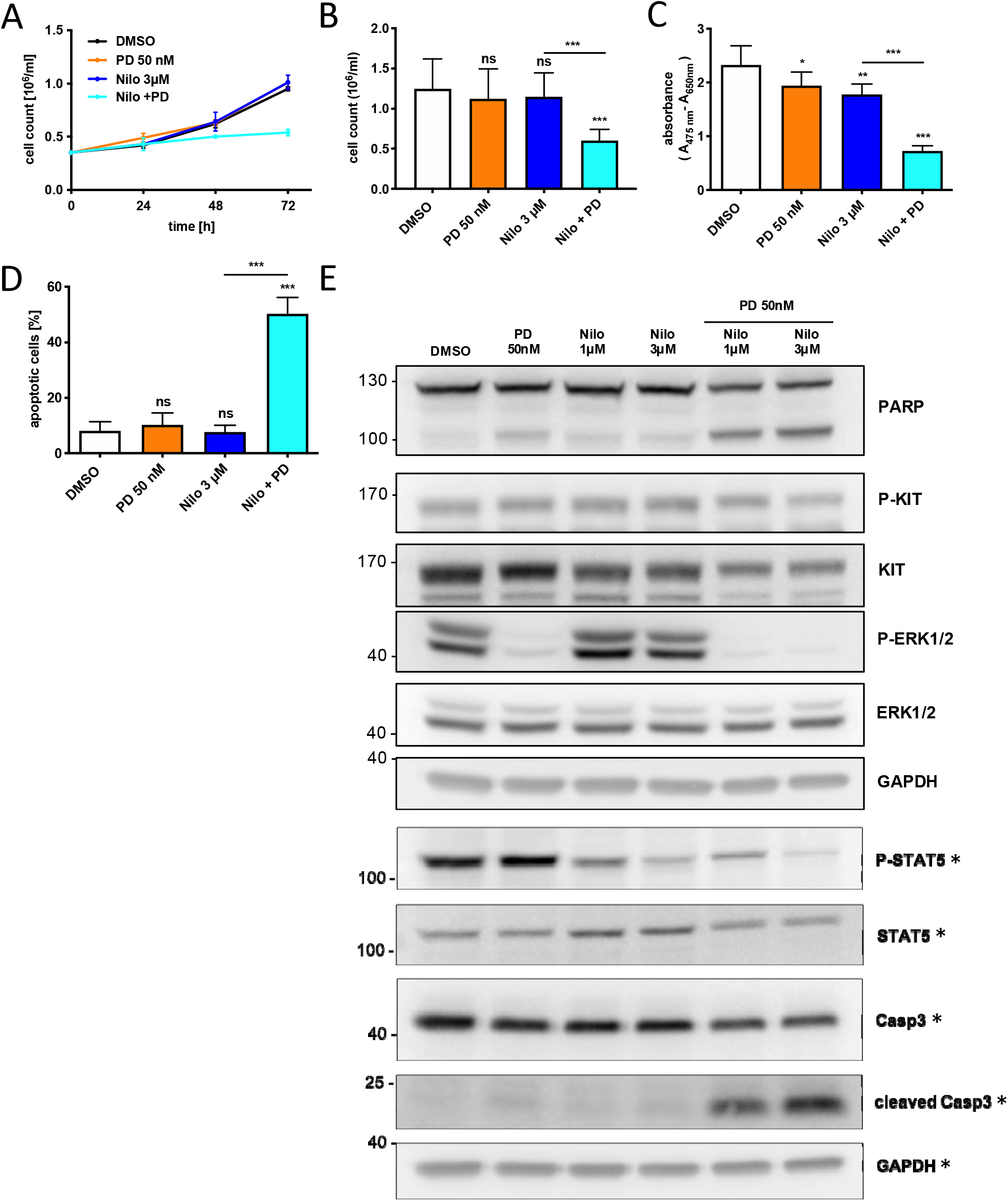
TKI Nilotinib synergizes with low dose MEK inhibitor PD0325901 in HMC-1.2 cells. HMC-1.2 cells were seeded at a density of 3.5 x 10^5^ cells/ml and treated with the solvent control DMSO, Nilotinib (3µM), PD0325901 (50nM) or in combination. Cells were counted every 24 h for up to 72 h using a CASY Cell Counter. (A) shows a time course of an individual experiment and (B) the average of repetitive experiments (n=5) after 72 h. The metabolic activity of treated cells was analyzed after 72 h using XTT assays (C) (n=6). Cell viability of treated cells was measured after 72 h by Annexin V and Propidium Iodide staining (D) (n=3). Lysates of cells treated for 3 h were analyzed by western blot for phosphorylation of KIT, ERK1/2 and STAT5. Caspase-3 activity was analyzed by detection of fragments of PARP and Caspase-3. GAPDH, KIT, ERK1/2, STAT5 and Caspase-3 served as loading controls (E), representative experiments (n=2). Mean + SD, one-way ANOVA followed by Tukey (multiple comparison) post-test. * P < 0.05, ** P < 0.01, and *** P < 0.001; ns indicates no significance.

### 4. Trametinib and Ponatinib represent an effective tandem for the suppression of HMC-1.2 cell proliferation and survival

Next, we combined Trametinib (10 nM) with the TKI Nilotinib and measured their combined effects on HMC-1.2 cells. As with PD0325901 (50 nM), Trametinib (10 nM) together with Nilotinib (3 µM) lead to a significant, synergistic repression of proliferation, metabolic activity, and HMC-1.2 survival (Fig. 4 A-C). Finally, we combined Trametinib (10 nM) with the third-generation TKI Ponatinib (300 nM), which we found to be more potent than the combination of Nilotinib with PD0325901 (by a factor of 10) concerning inhibition of HMC-1.2 cells (Fig. 3). This “low-dose” combination was observed to be very effective in the suppression of proliferation, metabolic activity, and survival of HMC-1.2 cells (Fig. 4 D-F). In conclusion, low concentrations of the TKI Ponatinib and the MEK inhibitor Trametinib effectively induce cell death of Imatinib-resistant HMC-1.2 cells by implementing synthetic lethality.

**Figure 4:**
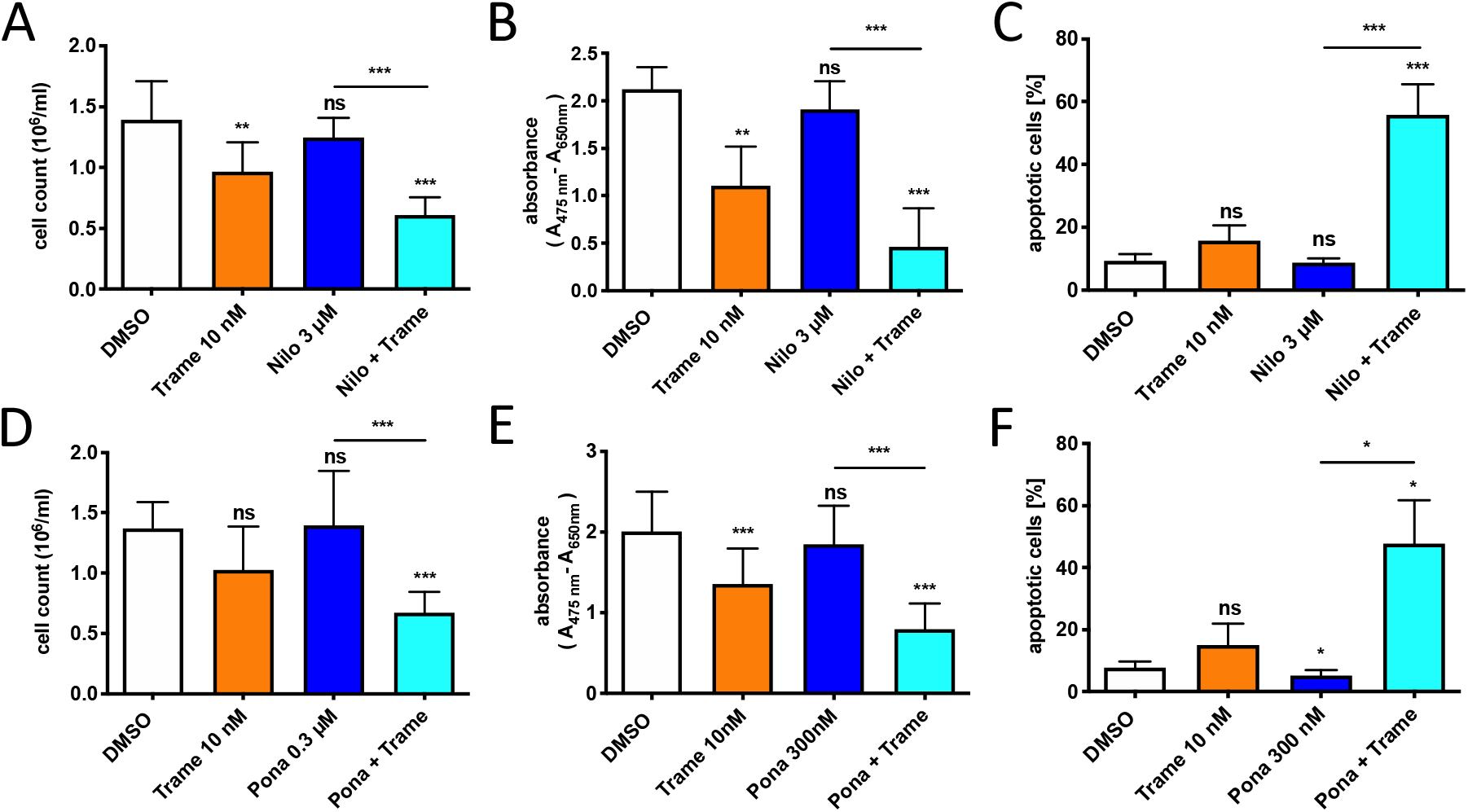
Next generation MEK inhibitor Trametinib has a higher potency in combination with the TKIs Nilotinib or Ponatinib in HMC-1.2 cells. HMC-1.2 cells were seeded at a density of 3.5 x 10^5^ cells/ml and treated with the solvent control DMSO, TKIs Nilotinib (3 µM) or Ponatinib (0.3 µM) alone or in combination with Trametinib (10 nM) for 72 h. Proliferation of Trametinib treated cells with Nilotinib (A) (n=4) or Ponatinib (D) (=5) was measured using a Casy cell counter. Metabolic activity of Trametinib treated cells with Nilotinib (B) (n=4) or Ponatinib (E) (n=5) was analyzed by XTT assays and survival of Trametinib treated cells with Nilotinib (C) (n=3) or Ponatinib (n=4) (F) was measured by Annexin V and Propidium Iodide staining. Mean + SD, one-way ANOVA followed by Tukey (multiple comparison) post-test. * P < 0.05, ** P < 0.01, and *** P < 0.001; ns indicates no significance.

### 5. Synergistic inhibition of proliferation and survival in ROSA^KIT D816V^ cells by the combination of Ponatinib and Trametinib

Though HMC-1.2 cells have been a valuable tool for investigating the molecular role and inhibitor susceptibility of KIT D816V in MCL, they exhibit certain weaknesses: a) HMC-1.2 cells most likely expressed additional mutations already when these MCL cells were isolated from an MCL patient and have acquired further potentially growth-promoting mutations since then; and b) HMC-1.2 cells express KIT V560G and D816V and not only KIT D816V and thus interference with the V560G mutation in the juxtamembrane region cannot be excluded. Therefore, we aimed at corroborating the synergistic effects of the Ponatinib/Trametinib combination in another cell line and made use of the SCF-independent, FcεRI-positive human MC line, ROSA^KIT D816V^, which has been generated from a stable stem cell factor (SCF) -dependent human MC line, ROSA (KIT WT), that has been transfected with *KIT* D816V (Saleh et al. 2014). To begin with, we titrated both inhibitors independently and measured their suppressive effects on ROSA^KIT D816V^ proliferation, metabolic activity, and survival. These cellular functions were attenuated by both inhibitors in a concentration-dependent manner (Fig. 5 A-F). Moreover, ROSA^KIT D816V^ cells appeared to be more sensitive to these inhibitors than HMC-1.2 cells (approximately by a factor of 10; compare Figs. 5 A-F to Figs. 1 C, F, I and 2 G, H, I). The combination treatment (Ponatinib, 100 nM; Trametinib, 1 nM) also resulted in a strong, synergistic suppression of ROSA^KIT D816V^ proliferation, metabolic activity, and survival Fig. 5 G-I). Compared to HMC-1.2 cells (Fig. 1 C), low concentrations of Ponatinib did not cause stronger proliferation in ROSA^KIT D816V^ cells (Fig. 5 A), suggesting that paradoxical activation of the MAPK pathway is not taking place in these cells. Indeed, whereas Ponatinib-induced CRAF/BRAF dimerization was detectable, increased ERK1/2 phosphorylation could not be observed (Fig. 5J), suggesting a more subtle effect than in HMC-1.2 cells. A comparable pattern was observed with the TKI Nilotinib (Fig. 5J), excluding a Ponatinib-selective effect. Nevertheless, the combination of low concentrations of Ponatinib and Trametinib synergistically inhibited proliferation and survival in MCs positive for KIT D816V, indicating that the presence of additional mutations in HMC-1.2 cells are not necessary for the successful treatment by this combination of inhibitors.

**Figure 5:**
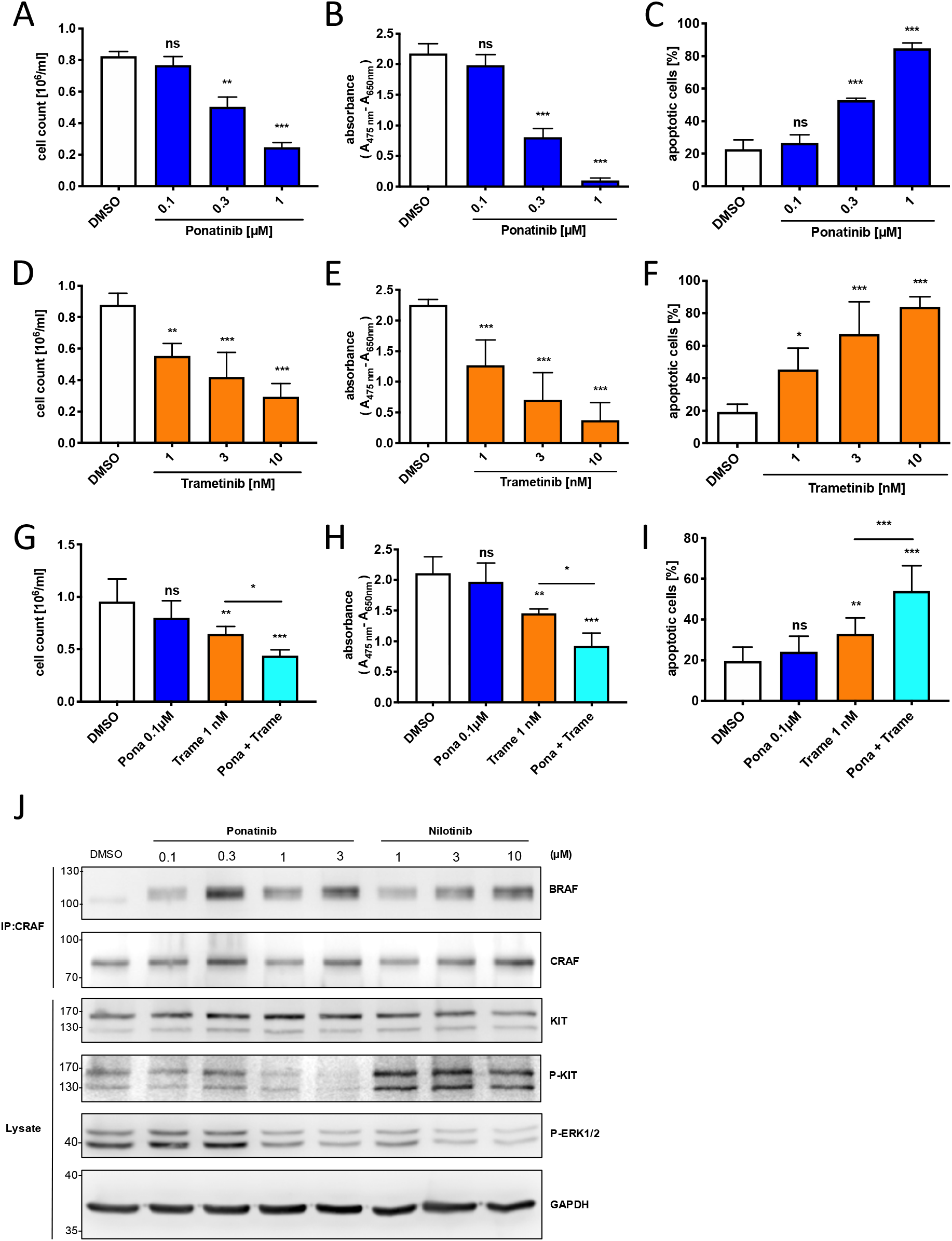
Combination of Ponatinib and Trametinib reduces proliferation and survival of Rosa ^KIT D816V^ cells. Proliferation (A), metabolic activity (B) or survival (C) of Ponatinib treated cells was analyzed after 72 h (n=3). Proliferation (D) (n=4), metabolic activity (E) (n=4) and survival (F) (n=3) of Trametinib treated cells was analyzed after 72 h. Proliferation (G) (n=4), metabolic activity (H) (n=3) and survival (I) (n=4) of single inhibitor (Ponatinib, Trametinib) treated cells or in combination was analyzed after 72 h. CRAF immunoprecipitations and total cell lysates from TKI (Ponatinib, Nilotinib) treated cells for 3 h were analyzed by western blot for CRAF, BRAF and phospho-KIT, KIT (loading control), phosphor-ERK1/2, GAPDH (loading control) (J), representative experiment (n=3). Mean + SD, one-way ANOVA followed by Tukey (multiple comparison) post-test. * P < 0.05, ** P < 0.01, and *** P < 0.001; ns indicates no significance.

### 6. MEK inhibition promotes TKI action on patient-derived iPS cells

The data of the previous sections support the idea that the addition of a MEK inhibitor to TKI-treated KIT D816V-positive cells promotes their inhibitory potential by preventing the impact of paradoxical RAF activation. SM and MCL are rare and heterogenous diseases. Although the majority of respective MCs are positive for *KIT* D816V, additional mutations might determine the severity and quality of the disease by affecting the pathogenic molecular signaling and response to therapy. Based on our results with the *KIT* D816V-positive cell lines HMC-1.2 and Rosa ^KIT D816V^, the aim was to verify our findings in a cellular model genetically closer to the patient situation. For this purpose, SM patient-derived iPS cells were used, which were previously established as a model system to test potential personalized therapies *in vitro* (Toledo et al. 2020). We tested *KIT* WT and *KIT* D816V iPS cell lines derived from two SM patients that are described on Stem Cell Registry (www.hpscreg.eu). Here, NGS analysis revealed additional single nucleotide polymorphisms (SNPs) such as of ASXL1, which has been reported for SM and other myeloid malignancies (Jawhar et al. 2015). Importantly, no clinical implications have been reported for those mutations. In contrast to this, patient 1 control iPS cells were positive for a TET2 ( p.Cys973Alafs*34) as well as a NRAS (p.Gly12Asp) mutations with likely pathogenic significance.

*KIT* WT and *KIT* D816V iPS cells were differentiated towards hematopoietic progenitors and myeloid cells, which was verified by detection of myeloid progenitors (Mp), granulocytes (Gr) and macrophages (Mac) (Supplementary Figure 1A). iPS cell-derived hematopoietic cells (Supplementary Figure 1B) were enriched for KIT expressing cells by MACS and KIT+ cells were used for inhibitor treatment. After 72 h of inhibitor treatment, XTT assays were performed to measure the impact on the metabolic activity of the cells. Consistent with and corroborating our recent experiments in HMC-1.2 and ROSA ^KIT D816V^ cells, a stronger decrease in metabolic activity was observed when Ponatinib was combined with Trametinib in all tested patient-derived cell lines independent of their *KIT* status and their individual genetic profiles (Fig. 6A, B).

**Figure 6:**
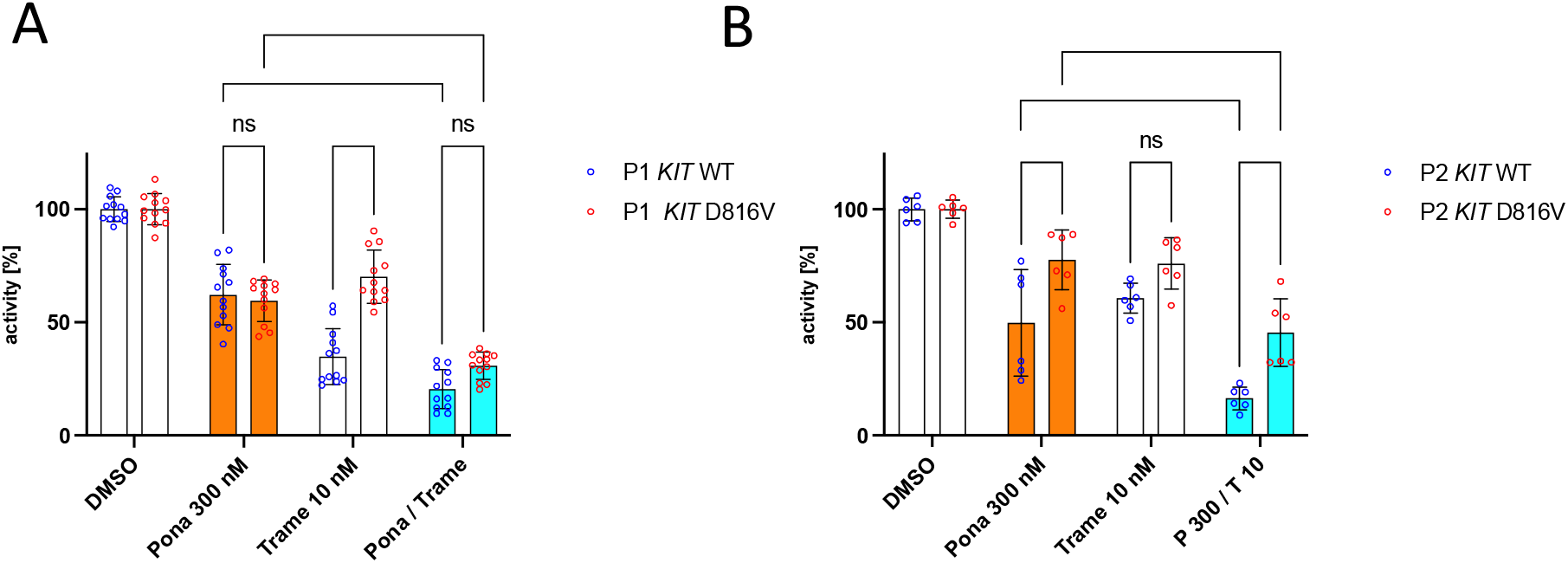
MEK inhibition promotes TKI action on patient derived iPS cells. *KIT* D816V and control *KIT* WT iPS cell-derived hematopoietic cells from 2 patients were incubated with the indicated inhibitor concentration as a single treatment or in combination. DMSO was used as solvent control. Metabolic activity was measured after 72h by XTT assays. Relative metabolic activity ± SD of KIT D816V and control iPS cell-derived *KIT* WT cells of (A) Patient 1 (n=4) and (B) patient 2 (n=2). Single dots show data of technical replicates. Mean ± SD, Two-way ANOVA followed by Tukey (multiple comparison) post-test. * P < 0.05, ** P < 0.01, and *** P < 0.001; ns indicates no significance.

Single treatments with Ponatinib reduced the activity of both patient 1 (P1)-derived cell lines carrying an additional *ABL1* mutation (Ser991Leu) of mutation class 2 (likely not pathogenic or of little clinical significance). This was expected since Ponatinib, due to its broad inhibitory range (O’Hare et al. 2009), is also able to affect KIT WT-positive cells (Fig. 6A). Interestingly, P1-derived *KIT* WT cells did respond stronger to single Trametinib treatment than *KIT* D816V cells. In agreement, genomic analyses of P1-derived KIT^WT^ iPS cells could identify an *NRAS* G12D mutation, which was not found in P1-derived *KIT* D816V-positive cells, most likely accounting for the strong response to MEK inhibition (Fig. 6A). Hematopoietic cells of a second patient (P2) did also respond to single Ponatinib treatment (Fig. 6B). The response to Ponatinib and the inhibitor combination was noticeably stronger in *KIT* WT cells. Nevertheless, combined inhibition of P2 *KIT* D816V cells by Ponatinib and Trametinib resulted in evident suppression of metabolic activity compared to these inhibitors alone (Fig. 6B). In conclusion, the analysis of *KIT* D816V cell lines together with patient-specific iPS cell-derived hematopoietic progenitors could confirm the benefit of MEK inhibition in addition to TKI treatments by suppressing the effect of paradoxical RAF activation. Moreover, taking into account the inhibitory profile of the applied TKI as well as thorough genetic analyses of patient samples are mandatory for a successful personalized therapy.

## Discussion

Our present study demonstrates that a combination of TKIs and MEK inhibitors is able to significantly increase the efficacy of anti-leukemia treatment compared to the single-drug regimens in Imatinib-resistant cells. Not only was the combined treatment more efficient in terms of strength of the anti-proliferative and pro-apoptotic effects, but it was also able to reach the desired beneficial effects at significantly reduced inhibitor concentrations. The possibility of applying lower drug doses suggests a successful reduction of unwanted detrimental side effects.

To date, TKIs are the gold standard for the treatment of proliferative diseases such as MCL and CML, caused and/or promoted by constitutively active tyrosine kinases. Unfortunately, mutants of respective tyrosine kinases, which are completely resistant to TKIs of the first generation, such as Imatinib, and largely resistant to second generation TKIs, such as Nilotinib, exist. These mutants comprise, for instance, KIT D816V and BCR-ABL1 T315I. Particularly KIT D816V poses a significant problem since more than 80% of patients suffering from various forms of SM express this constitutively active mutant of KIT in their aberrant MCs (Valent et al. 2017). HMC-1.2 MCL cells expressing KIT V560G and D816V have been used to identify TKIs, which are able to reduce or even prevent KIT D816V kinase activity, such as Nilotinib (AMN107; IC_50_≍2363nM), Midostaurin (PKC412; IC_50_≍191nM) (Gleixner 2006) as well as Ponatinib (IC_50_ between 0.05 - 0.5 µM) (Gleixner et al. 2013). Most of these TKIs have a broad target profile. Midostaurin, for instance, has been identified as an inhibitor of multiple tyrosine kinases (e.g. SYK, FLK1, KIT, FGR, SRC, FLT3, PDGFRβ, and VEGFR1/2) as well as serine/threonine kinases (e.g. PKC-α/β/γ, AKT, and PKA) with IC_50_ values ranging from 80-500 nM (Peter et al. 2016). Ponatinib also targets various tyrosine kinases like KIT, ABL, PDGFRα, VEGFR2, FGFR1, and SRC (Tan et al. 2019). Tyrosine phosphorylation profiling and/or chemical proteomics for TKIs as for example Imatinib, Nilotinib, and Dasatinib in BCR-ABL-positive CML cells but also other cancer cell lines documented this variety of targets for single TKIs (Preisinger et al. 2013; Giansanti et al. 2014; Rix et al. 2007). Unexpectedly, also binding to and inhibition of a non-kinase protein, the oxidoreductase NQO2, was demonstrated (Rix et al. 2007) expanding the quality of TKI target proteins and pointing even more to the necessity of reducing drug concentrations, thereby prohibiting the occurrence of severe side effects due to unwanted inhibition of alternative targets.

An additional drawback of some TKIs is their binding to and induction of dimer formation of RAF kinases enabling their activation by RAS-GTP, a phenomenon called “paradoxical activation”. Packer et al. have pointed out that a combination of the TKI Nilotinib with MEK inhibitors induces synthetic lethality of CML cells, thereby preventing the consequences of paradoxical RAF activation, namely enhanced proliferation and survival (Packer et al. 2011). Importantly, the phenomenon of paradoxical RAF activation is strongly dependent on the presence of active RAS, which can be provided directly by activating RAS mutations or indirectly by active upstream signalling elements, such as KIT D816V or BCR-ABL1 T315I.

While TKIs bind to similar structured ATP binding sites, MEK inhibitors bind to a unique inhibitor-binding pocket that is separate from but adjacent to the Mg^2+^-ATP-binding site in MEK1 and MEK2 (Ohren et al. 2004). The benefit of using the feedback buster Trametinib was the prevention of accumulation of phosphorylated MEK and a higher efficiency compared to the classical MEK inhibitor PD0325901. Trametinib is used in a number of clinical trials (197 – reference date 08/2020; source clinicaltrials.org), 9 studies are in phase 3 and three studies are in phase 4 (melanoma, non-small cell lung cancer, various other solid tumours and astrocytoma).

Numerous studies have shown that the MEK/ERK pathway is important for proliferation and survival of tumour cells. In this line, in HMC-1.2 cells, PD0325901-mediated MEK inhibition resulted in significant reduction of *BCL2L1* as well as *CCND1* expression (Fig. 2). Cyclin D1 (CCND1) is the regulatory component of the CCND1-CDK4 complex that phosphorylates and inhibits RB1 and thereby allows the cell to proceed through the G_1_/S phase of the cell cycle (Matsushime et al. 1992). The RAF-MEK-ERK1/2 pathway was shown to be important for G_1_/S cell cycle progression by the positive regulation of *CCND1* expression (Lavoie, Rivard, et al. 1996; Lavoie, L’Allemain, et al. 1996). In addition to proliferation, MEK/ERK signalling also promotes survival, for instance of human pancreatic cancer cells, by regulating the expression of anti-apoptotic BCL2 family members (Boucher et al. 2000). Moreover, ERK1/2 activation leads to repression of pro-apoptotic *BCL2L11* expression (Weston et al. 2003), and can promote dissociation of BIM-EL from BCL2 family members (Ewings et al. 2007), hence impeding apoptosis.

In both *KIT* D816V-positive MC lines studied (HMC-1.2 and ROSA^KIT D816V^), different TKIs induced paradoxical activation manifesting in BRAF-CRAF dimerization and ERK1/2 activation, which enabled repression of proliferation and promotion of apoptosis by combinations of low concentrations of the used TKIs and MEK inhibitors. Nevertheless, differential effects were monitored in the presence of TKIs only. While in HMC-1.2 cells low TKI concentrations induced a significant increment in proliferation, this was not the case in TKI-treated ROSA^KIT D816V^ cells, indicating additional pro-proliferative signalling processes in HMC-1.2 cells. The patient-derived HMC-1.2 cells carry an additional mutation in the juxtamembrane region of KIT (V560G) (Butterfield et al. 1988) and most likely additional mutations frequently detected in MCL cells (e.g. in *ASXL1*, *SRSF1*, or *TET2*) (Valent et al. 2017). ROSA^KIT D816V^ cells, however, were generated by lentiviral transduction with *KIT* D816V of the human umbilical cord blood-derived MC line ROSA^KIT WT^ (Saleh et al. 2014). Further differences pertain to the respective culture conditions. Whereas HMC-1.2 cells were maintained in culture medium only containing foetal bovine serum, ROSA^KIT D816V^ cells were grown in medium containing additional nutrients and supplements (e.g. insulin/transferrin, vitamins, and nucleotides). Though the exact reason for this difference in TKI-induced proliferation is not yet clear, this discrepancy indicates that paradoxical activation of the MAPK pathway is not coupled to increased proliferation in a mandatory manner.

Another question remains concerning the apparent inability of the TKI Imatinib to cause a proliferative response in HMC-1.2 cells despite its ability to induce paradoxical activation of the MAPK pathway. This was also observed in various cell lines (D04, SW620, H460, Panc1, K562, Ba/F3-BCR-ABL^T315I^) by Packer et al. (Packer et al. 2011). As referred to above (Preisinger et al. 2013; Giansanti et al. 2014; Rix et al. 2007), TKIs strongly interact with a variety of different kinases (even with non-kinases); hence in contrast to Nilotinib and Ponatinib, Imatinib might cause signals that counteract the positive effect of the paradoxical activation even in the presence of active ERK1/2. Moreover, as mentioned for ROSA^KIT D816V^ cells, the coupling of paradoxical activation of RAF kinases to a proliferative response appears to be not mandatory.

Recent advances in the generation of patient-derived iPS cells together with next generation sequencing (NGS) has been established to characterize patient-specific cancer cell profiles in order to facilitate personalized therapy. Although our data could reflect the heterogeneity of MCL between the patient-derived iPS cell lines generated from two patients, the identified inhibitor combination was continuously effective. In summary, the combination of the MEK inhibitor Trametinib with the TKI Ponatinib induced a synergistic inhibition of proliferation and survival in *KIT* D816V-positive model systems. Moreover, the reduction of inhibitor concentrations achieved by the combination of MEK inhibitor and TKI could increase the safety and tolerability of the anti-leukemia therapy. This renders the combinatorial treatment a reliable and powerful approach and might be a beneficial weapon to fight leukemic cells.

## Acknowledgements

We are thankful to S. Capellmann and Dr. C. Preisinger for proofreading and discussions.

---

**Supplemental Figure 1:**
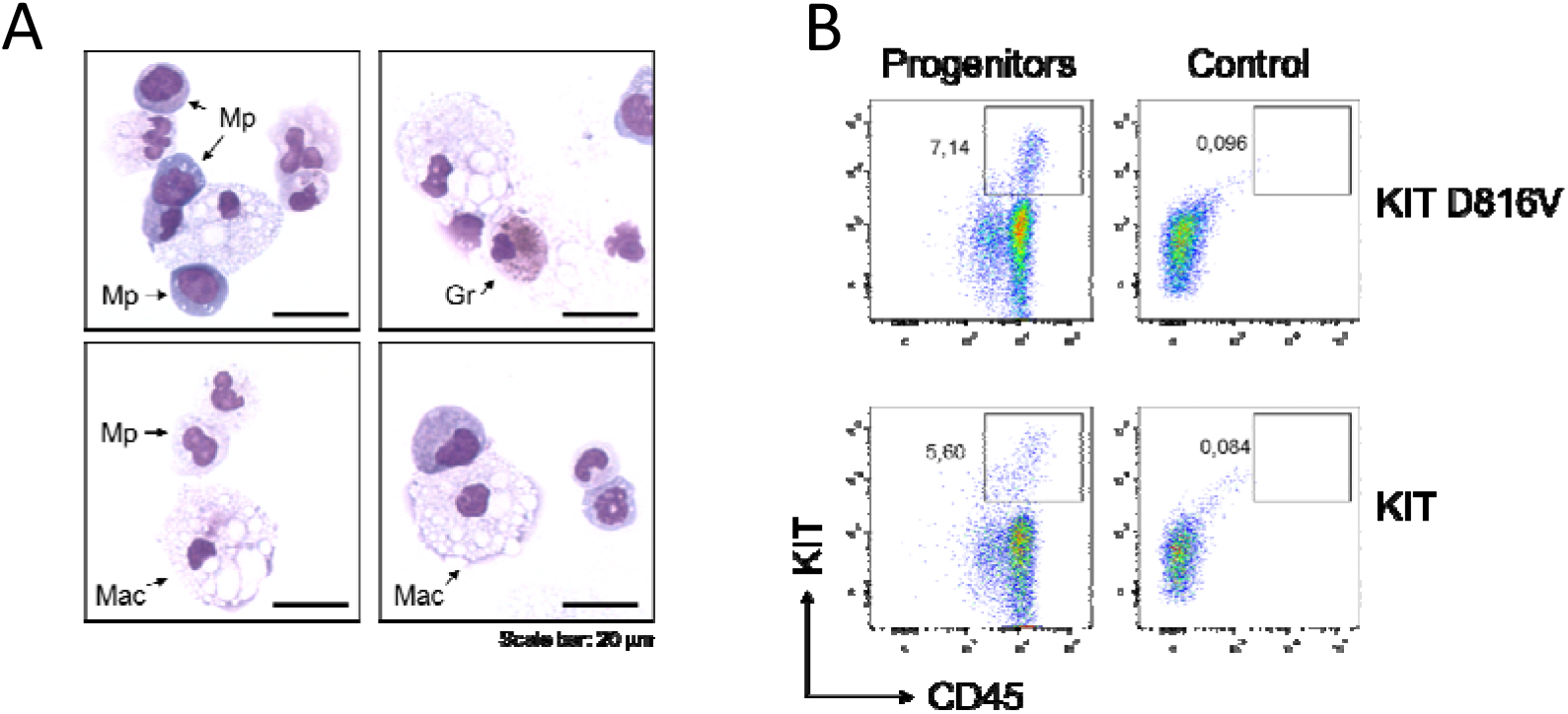
Generation of *KIT* D816V and *KIT* WT iPS cell-derived hematopoietic cells. (A) Representative light microscopy images of Diff-Quick stained cytospin preparations of iPS cell-derived hematopoietic cells. Mp: myeloid progenitors. Gr: granulocytes. Mac: macrophages. Scale bar: 200 µm (B) Representative flow cytometry dot plots of iPS cell-derived hematopoietic cells harboring the *KIT* D816V mutation (top) or wild-type KIT (bottom) showing CD45^+^KIT^+^ population.

